# Genetically encoded photocatalytic protein labeling enables spatially-resolved profiling of intracellular proteome

**DOI:** 10.1101/2022.08.01.502286

**Authors:** Fu Zheng, Chenxin Yu, Xinyue Zhou, Peng Zou

## Abstract

Mapping the subcellular organization of proteins is crucial for understanding their biological functions. Herein, we report a reactive oxygen species induced protein labeling and identification (RinID) method for profiling subcellular proteome in the context of living cells. Our method capitalizes on a genetically encoded photocatalyst, miniSOG, to locally generate singlet oxygen that reacts with proximal proteins. Labeled proteins are conjugated *in situ* with an exogenously supplied nucleophilic probe, which serves as a functional handle for subsequent affinity enrichment and mass spectrometry-based protein identification. From a panel of nucleophilic compounds, we identify biotin-conjugated aniline and propargyl amine as highly reactive probes. As a demonstration of the spatial specificity and depth of coverage in mammalian cells, we apply RinID in the mitochondrial matrix, capturing 394 mitochondrial proteins with 97% specificity. We further demonstrate the broad applicability of RinID in various subcellular compartments, including the nucleus and the endoplasmic reticulum.

## INTRODUCTION

Within highly compartmentalized eukaryotic cells, the subcellular localization of proteins is crucially linked to their biological functions. This is most readily observed in secreted proteins, which constantly traffic through the endoplasmic reticulum (ER)-Golgi apparatus-plasma membrane axis^1^. In signal transduction pathways or stress response pathways, activated protein factors often translocate from cytoplasm^2^ or mitochondria^3^ into the nucleus to initiate the transcription of effector genes. Thus, our understanding of protein function would require knowledge of not only the abundances and activities of individual proteins, but also their spatial arrangements, ideally at the proteome level and in the native cellular context^4^.

To profile the subcellular organization of proteome, a number of spatial-specific chemical labeling techniques have been developed over the past decade, which often capitalize on genetically targetable enzymes (e.g. APEX^5, 6^, BioID^7^, TurboID^8^) that catalyze the formation of short-lived and highly reactive intermediates in live cells (e.g. biotin-conjugated phenoxyl radicals^5,6^ or biotinyl 5’-adenylate^7, 8^). These intermediates react with proteins in close proximity to their source of generation, thus achieving high spatial specificity of labeling. Proximity labeling enzymes have been applied to investigate the proteome of many subcellular compartments, including the mitochondria^9^, endoplasmic reticulum^10^, primary cilia^11^, etc.

However, the requirement of using H2O2 in APEX labeling may cause toxicity to living samples^12^. Cellular expression of constitutively active TurboID would lead to high background biotinylation of endogenous proteins, which may interfere with cellular physiology and cause cytotoxicity. These problems have motived us to search for a new proximity labeling method, where the activation avoids toxic H2O2 and could be controlled by light trigger, to minimize the impact on cell physiology.

Herein, we report a light-activatable proximity labeling method with minute-level turn-on kinetics, excellent labeling efficiencies, and high spatial specificity in various organelles. A protein photocatalyst, miniSOG, is genetically targeted to various subcellular compartments, where it generates singlet oxygen upon blue light illumination^13^. Reactive and short-lived singlet oxygen then reacts with proteins in close proximity to miniSOG, thus offering high spatial specificity of labeling. Photo-oxidized protein intermediates are intercepted with a nucleophilic probe and subsequently enriched for mass spectrometry analysis. We screened a panel of nucleophilic compounds and identified biotin-conjugated aniline and propargylamine as highly reactive probes. Application of RinID in the mitochondrial matrix identifies 394 proteins with 97% mitochondrial specificity, which compares favorably to previously reported methods. RinID can also be applied in other subcellular compartments (e.g. nucleus and ER), thus demonstrating its broad applicability and capability of complementing other proximity labeling methods.

## RESULTS

Proteins are prone to be oxidized by reactive oxygen species (ROS)^14^. For example, the imidazole ring of histidine could be oxidized into 2-oxo-imidazole in the presence of singlet oxygen^15^. In cells, extensive oxidation of a protein could hamper its enzymatic activity or interaction with other biomolecules, thus turning off its function. This feature has been leveraged in chromophore-assisted light inactivation (CALI^16^) strategy to achieve selective photo-ablation of specific proteins in live cells. miniSOG, an engineered flavin-binding protein, has been employed in CALI experiments as a protein fusion tag, which generates singlet oxygen upon blue light illumination. In this study, we aim to repurpose miniSOG for ROS-induced proximity-dependent proteome labeling and identification (RinID). We propose to capture the protein photo-oxidation intermediates *in situ* with amine-based nucleophilic probes functionalized with an affinity purification handle. We reason that, due to the short lifetime (<0.6 μs) and limited diffusion radius (∼70 nm) of singlet oxygen^17^, such labeling reaction occurs only proximal to miniSOG, and the labeled proteins could be subsequently enriched and identified through mass spectrometry-based proteomic analysis (Figure 1A).

**Figure 1.**
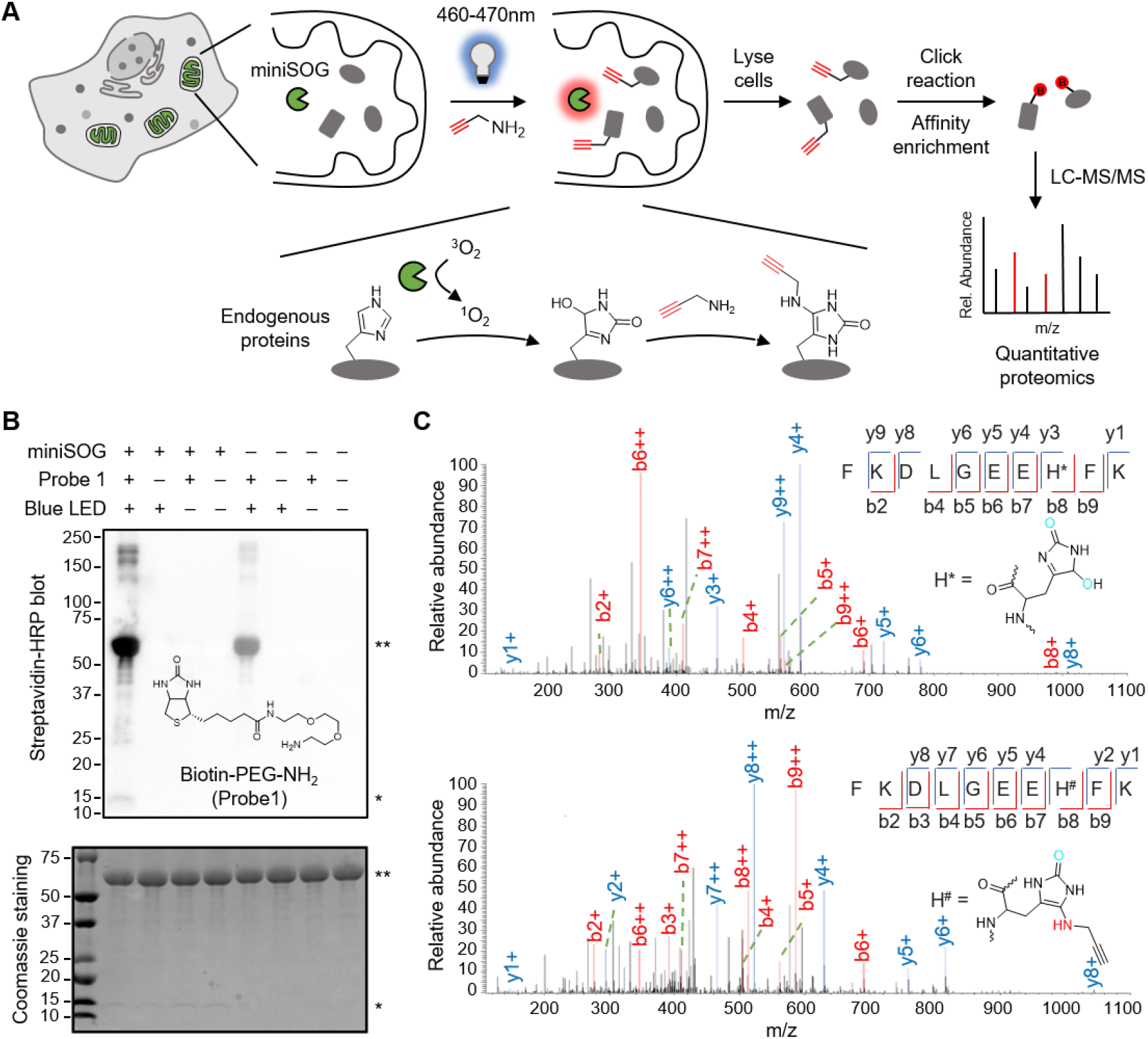
Experimental scheme and *in vitro* characterization of ROS-induced proximity-dependent protein labeling and identification (RinID). **A)** Scheme of RinID workflow. **B)** Western blot and SDS-PAGE analysis of miniSOG-mediated photo-oxidation of the model protein, bovine serum albumin (BSA) with 20mM Biotin-PEG-NH2 probe. Samples were illuminated with blue LED at 19 mW·cm^-2^ for 30 min. * miniSOG, ** BSA.**C)** MS/MS spectra of a representative peptide with histidine residue oxidized to 5-hydroxy-1,5-dihydro-2-oxoimidazole (top) and 5-propargylamino-1,5-dihydro-2-oxoimidazole (bottom).

We started by testing miniSOG-mediated photo-oxidation *in vitro* with a model protein, bovine serum albumin (BSA, PDB: 4F5S), with biotin-conjugated alkyl amine (biotin-PEG-NH2, probe **1**) as the nucleophilic probe (Figure 1B, S1A-B). In the presence of 100 μM purified miniSOG and 20 mM biotin-PEG-NH2, BSA in phosphate buffer saline solution (pH 7.3) was illuminated with 460-470 nm blue LED at the mild intensity of 19 mW·cm^-2^ for 30min at room temperature. Western blot analysis showed successful biotinylation of BSA (Figure 1B). In negative controls omitting either miniSOG or light illumination, the biotinylation signal was substantially reduced. We attributed the low biotinylation background in the absence of miniSOG to the residual serum-derived photosensitizer impurities in the BSA sample. We repeated the labeling with another nucleophilic probe, propargyl amine (PA) and obtained similar results (Figure S1C). Together, the above characterizations demonstrate that miniSOG is capable of labeling proteins with amine-conjugated probes in a blue light-dependent manner.

To understand the labeling mechanism, we searched for the amino acid residues of photo-oxidation and PA conjugation by mass spectrometry. Photo-oxidized BSA sample was proteolytically digested into peptide fragments and analyzed by liquid chromatography-tandem mass spectrometry (LC-MS/MS). Among the five amino acid residues (histidine, tyrosine, tryptophan, cysteine, and methionine) that are commonly oxidized by singlet oxygen^14^, the photo-oxidation products of histidine were most readily detected on the mass spectrometry, with observed mass shifts of 31.990 Da and 69.022 Da matching the transformation of imidazole ring into 5-hydroxy-1,5-dihydro-2-oxoimidazole and 5-propargylamino-1,5-dihydro-2-oxoimidazole (Figure 1C, S1D). This observation is consistent with a recent report that amine probe 1-methyl-4-arylurazole could react with photocatalytically oxidized histidine residue^18^. In both cases, singlet oxygen reacts with the imidazole ring to form an endoperoxide intermediate, which undergoes nucleophilic addition at the C4 position by either water or an amine probe, generating C-O and C-N bond, respectively. Consistent with this mechanism, the m/z of both products were identified at solvent-exposed His18 and His378 sites of BSA following blue LED irradiation in the presence of PA (Figure 1C). In addition, although we have also identified the oxidized products of tryptophan, tyrosine, and methionine, we have failed to detect their nucleophilic addition products with PA (Table S1, Figure S1E-H). We conclude from the above data that miniSOG could mediate the photocatalytic protein conjugation with amine probes. Due to the competition from water and other nucleophilic species in living cells, probe labeling needs to be optimized for efficient protein capture.

We next sought to achieve miniSOG-mediated protein labeling in live cells. We constructed human embryonic kidney 293T (HEK293T) and HeLa cell lines targeting miniSOG to various subcellular compartments, including both membrane-bound (e.g. mitochondrial matrix, ER) and membraneless organelles (e.g. stress granule) (Figure 2A). Meanwhile, we prepared a panel of biotin-conjugated amine probes, including primary alkyl amine, aniline, naphthylamine, and hydrazide, which differ in nucleophilicity, steric hindrance and basicity (Figure 2B, S2A). For comparison, we also included biotin-conjugated phenol, the commonly used substrate for APEX^5,6^. Following probe incubation at 0.5 – 2 mM in the culture medium for 5 min, cells were illuminated with blue LED at 43 mW·cm^-2^ for 15 min. Thereafter, cells were lysed and analyzed by Western blot. Among these probes, biotin-aniline (BA, probe 3) yielded the highest labeling signal (Figure S2B-D). We speculated that some of these probes may have limited permeability through the cell membrane, and re-evaluated their labeling efficiency in cell lysate. Notably, strong biotinylation signal was observed for all biotinylated probes in a miniSOG- and blue light-dependent manner (Figure S2E), thus indicating that low membrane permeability may contribute to the low labeling signal of biotin-conjugated probes in live cells.

**Figure 2.**
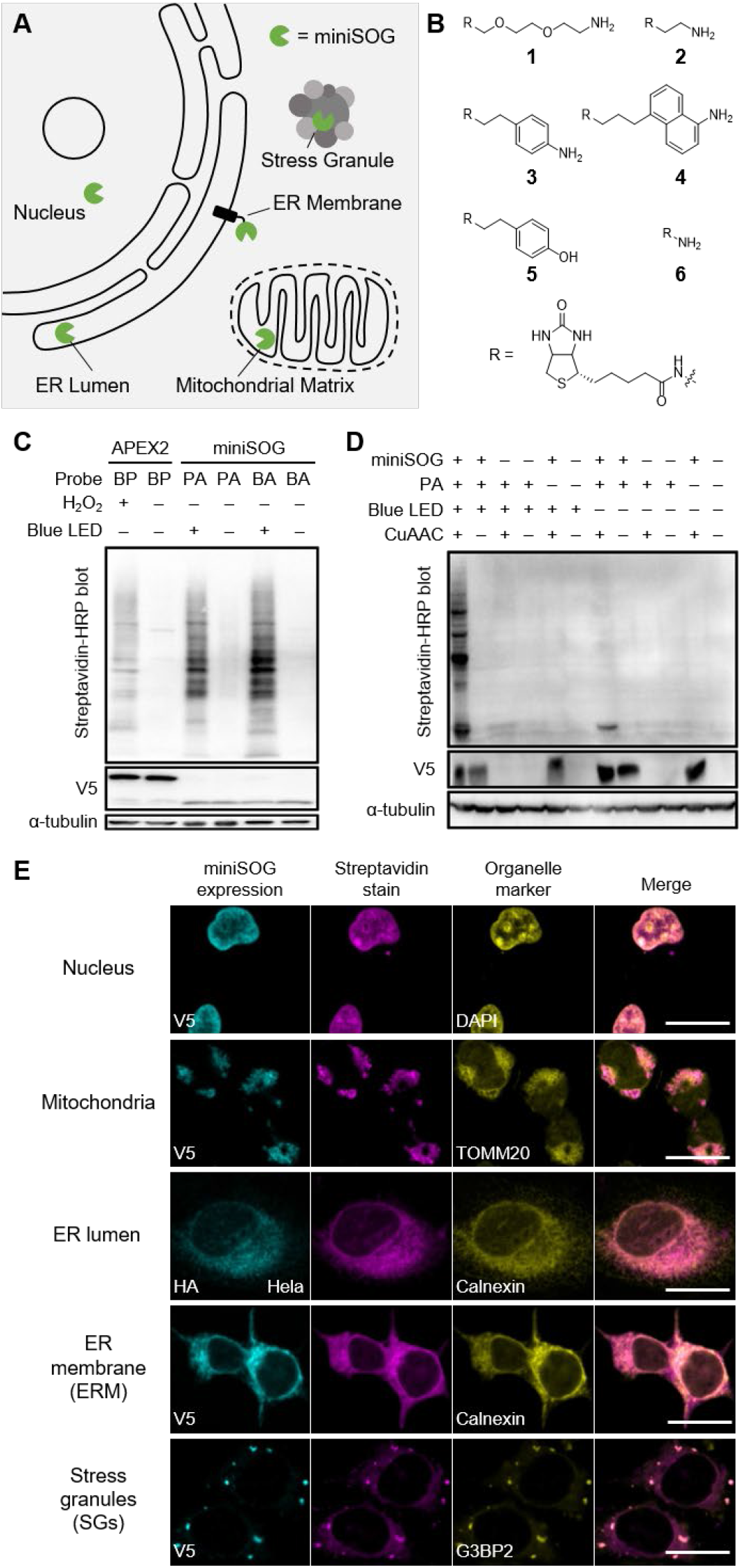
Application of RinID at different subcellular localizations in living cells. **A)** Subcellular targeting of miniSOG at different organelles. **B)** Chemical structures of probes used in this study. **C-D)** Western blot analysis comparing of mitochondrial matrix-targeted RinID labeling and APEX2 (C), and comparing RinID labeling against control experiments omitting (blue light illumination, PA probe, miniSOG, and/or CuAAC) (D). **E)** Confocal fluorescence images of cultured mammalian cells labeled with RinID in different organelles. Scale bar: 20μm. HEK293T cells were used unless otherwise noted.

To improve cell membrane permeability, we turn to the much smaller PA probe, whose excellent water solubility enables the probe to be supplied at higher concentration (e.g. 10 – 20 mM), which favorably competes with water and endogenous nucleophiles at intercepting the miniSOG-mediated protein photo-oxidation intermediate. Following miniSOG labeling and cell lysis, copper(I)-catalyzed alkyne-azide cycloaddition (CuAAC) click reaction was performed to install the biotin moiety (Figure 1A). Using mitochondrial matrix-targeted miniSOG as a model and Western blot signal intensity as the readout, we optimized PA probe concentration (Figure S3A) and light illumination time (Figure S3B), and determined 20 mM PA for 15 min as the optimal condition. Western blot analysis revealed that the overall biotinylation with 20 mM PA probe was comparable to 5 mM biotin-aniline probe, and higher than all other biotin-conjugated probes (Figure 2C and S2B-E). Notably, low levels of biotinylation background could be observed in negative controls omitting miniSOG or light illumination (Figure 2D), which we attributed to the presence of native photosensitizers (e.g. FMN) and naturally oxidized proteins in cells. To remove this background signal, ratiometric quantitative mass spectrometry experiments with proper controls should be performed.

We benchmarked the labeling efficiency of our method with APEX2, which has been widely used for proximity labeling. In the mitochondrial matrix, miniSOG-mediated protein labeling with PA and biotin-aniline are both substantially higher than APEX2-mediated labeling with biotin-phenol, even when miniSOG was expressed at a lower level than APEX2 (Figure 2C). However, it should be noted that miniSOG-mediate labeling typically requires light illumination for 15 min, whereas APEX2 requires only 1 min or less. Thus, when studying highly dynamic biological processes, such as G-protein coupled receptor signaling, APEX2 is still recommended for its fast reaction kinetics^19, 20^. Taken together, the above analysis established PA and biotin-aniline as suitable probes for RinID. We considered PA as a more cost-effective probe due to its commercial availability, which we used for all subsequent experiments.

To evaluate the spatial specificity of miniSOG-mediated protein labeling, we performed immunofluorescence imaging of cell samples labeled with PA (Figure 2E). Following photo-oxidation, cells were fixed and permeabilized with cold methanol. Biotinylation signal was detected by staining cells with streptavidin-conjugated fluorophores, while the localizations of miniSOG and the morphology of relevant organelles were visualized via antibody staining (or DAPI staining in the case of nucleus). Confocal fluorescence microscopy reveals good co-localization between biotinylated proteins and organelle markers, thus demonstrating the high spatial specificity of our method. In wild-type HEK293T cells lacking miniSOG and in negative controls omitting PA or light illumination, labeling was almost undetectable. However, we did notice the presence of a low biotinylation background that permeated throughout the cytoplasm and nucleus (Figure S4). This background was reminiscent of our previous observation in Western blot analysis, which was likely caused by endogenous photosensitizer and oxidized proteins. Collectively, RinID could label subcellular proteomes with high spatial specificity within 15 min at various membrane-bound and membraneless organelles in different cell lines.

We then evaluated the specificity and coverage of RinID with quantitative mass spectrometry (MS)-based proteomic profiling. HEK293T cells expressing mitochondrial matrix-targeted miniSOG were incubated with 20 mM PA and illuminated with blue LED at 30 mW·cm^-2^ for 15 min (Figure 3A). Following light illumination, cells were collected and lysed, and the lysate was reacted with biotin-conjugated azide via click reaction. Thereafter, biotinylated proteins were captured by streptavidin-coated agarose beads. Successful enrichment was confirmed by SDS-PAGE and silver staining (Figure S5).

**Figure 3.**
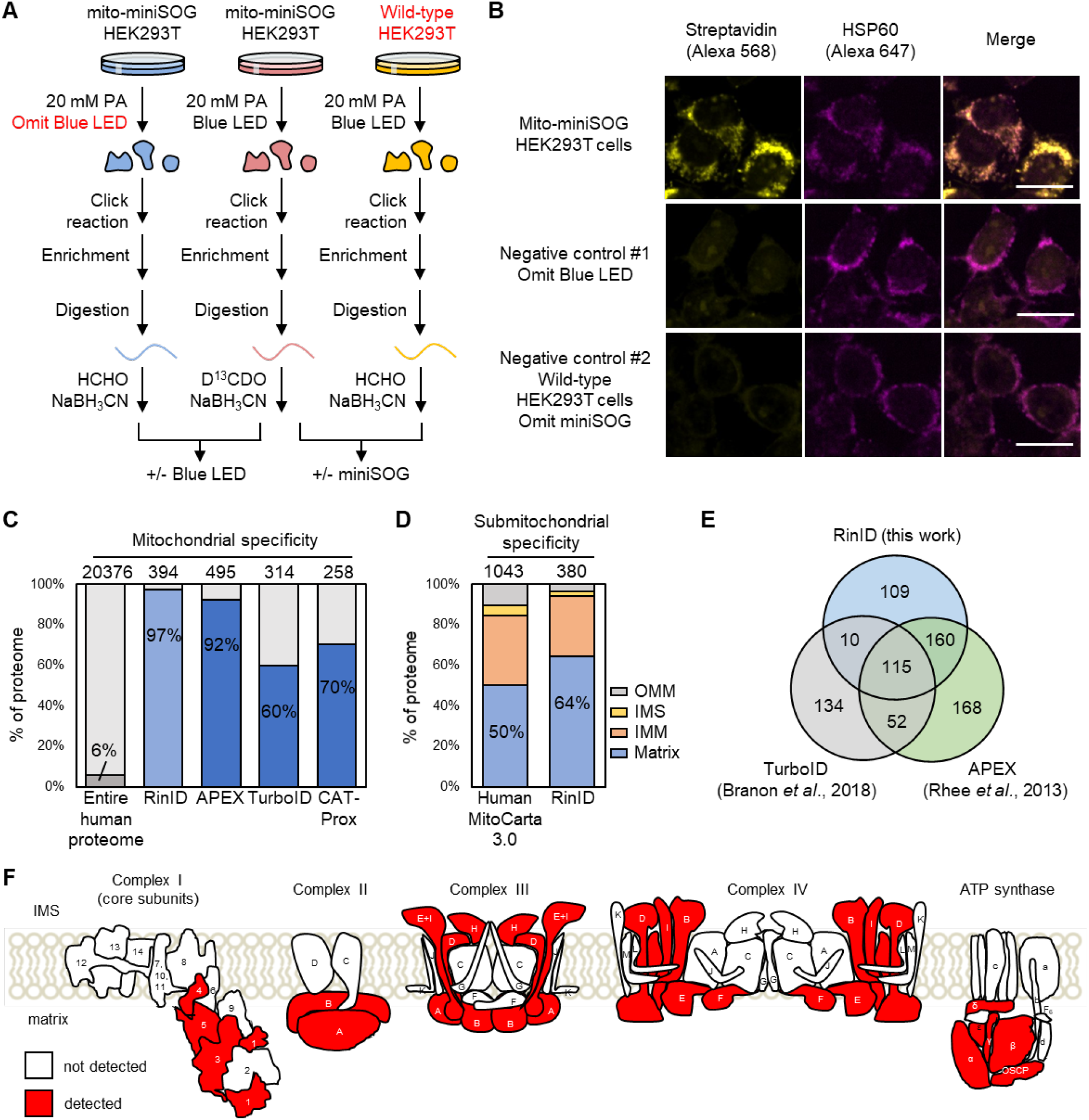
Analysis of mitochondrial proteome identified by RinID. **A)** Schematic workflow of mass spectrometry-based proteomic analysis of RinID at mitochondrial matrix. **B)** Confocal fluorescence images of HEK293T cells labeled with RinID and control samples omitting blue light or miniSOG. **C)** Comparison of spatial specificity of RinID proteomic data with three other proximity labeling methods at mitochondria, Mitocarta 3.0 are defined as mitochondrial proteome. **D)** Sub-mitochondrial specificity analysis for mitochondrial matrix RinID proteomic dataset. **E)** Comparison of mitochondrial proteome covered by RinID, APEX, and TurboID. **F)** Cartoon representation of RinID protein coverage in the electron transport chain complexes embedded in the mitochondrial inner membrane. miniSOG is targeted to the mitochondrial matrix side.

As mentioned above, background labeling should be carefully removed via quantitative MS experiments. For this purpose, we designed two negative controls and applied stable isotope dimethyl labeling strategy to quantitatively measure the ratios of protein abundance between samples. While one control omitted blue LED irradiation to account for PA labeling on native oxidized proteins, the other control used wild-type HEK293T cells lacking miniSOG to eliminate background protein labeling induced by endogenous photosensitizers (Figure 3A, 3B). For each set of experiments (+/-blue LED or +/- miniSOG), two biological replicates were performed. Both labeled samples and control samples went through the same enrichment workflow and subsequently digested by trypsin. The resulting peptides were treated with isotope-encoded formaldehyde (heavy D^13^CDO for labeled samples versus light HCHO for control samples) and NaBH3CN to methylate their –NH2 groups, leading to mass shifts of 34.0631 Da versus 28.0313 Da, respectively. The heavy and light samples were mixed and analyzed by LC-MS/MS for peptide identification and abundance determination.

A total of 1634 and 1882 proteins were identified and quantified in both replicates for “+/- blue LED” and “+/- miniSOG” datasets, respectively. For each dataset, proteins were ranked by their averaged H/L ratios and the cut-off ratios were determined with receiver operator curve (ROC) analysis (Figure S6, Table S2, Table S3). For ROC analysis, 878 proteins annotated as ‘mitochondrial matrix’ or ‘mitochondrial inner membrane’ in MitoCarta 3.0^21^, a well-established human mitochondrial proteome database, were defined as the ‘true positive’ list. The ‘false positive’ list was composed of 109 proteins with ‘mitochondrial outer membrane’ annotation in MitoCarta 3.0, as well as a manually curated list of 389 cytoplasmic proteins (Table S4). The cut-off H/L ratios were set at 2.95 and 2.93, yielding 486 and 495 enriched proteins for “+/- blue LED” and “+/- miniSOG” datasets, respectively. The overlap of these two lists contained 394 proteins, which was defined as our mitochondrial proteome (Table S2, S3, S5).

Notably, this protein inventory has exceptionally high mitochondrial specificity, with 97% (383 out of 394) of proteins listed in the MitoCarta 3.0, which is higher than previously reported proximity labeling methods, including APEX (92%)^5^, TurboID (60%)^8^, and small molecule photosensitizer-based CAT-Prox (70%)^22^ (Figure 3C). In terms of sub-mitochondrial specificity, 64% and 30% of our RinID dataset are mitochondrial matrix and inner membrane proteins, respectively (Figure 3D). To further examine the coverage and the spatial specificity of RinID, we use the electron-transport chain complexes as a model, whose membrane topology has been well resolved through structural biology studies. Figure 3F and Table S6 show that the majority of protein components identified by RinID are exposed to the mitochondrial matrix, where biotinylation occurs. The coverage of mitochondrial proteins by RinID is similar to that of APEX (495 proteins) and TurboID (314 proteins), with an overlap of 115 proteins, almost all of which (114 proteins) are annotated in the MitoCarta 3.0 database. In addition, 109 proteins are uniquely identified by RinID (Figure 3E), including 100 mitochondrial proteins (92%). This difference in coverage may arise from the preferences of three methods toward different amino acid residues: whereas APEX2 and TurboID favor tyrosine and lysine, respectively, RinID targets histidine. Taken together, the above comparisons indicate that RinID offers exceptional spatial specificity and good coverage, and could complement existing methods.

As a demonstration of its broad applicability, we further extended RinID to profile the local proteomes at two other subcellular compartments: the nucleus and the ER membrane (ERM). For the profiling of nuclear proteome, HEK293T cells expressing histone protein H2B-fused miniSOG were labeled by 10 mM or 20 mM PA for 15 min, in two replicated experiments. For the negative control, blue light illumination was omitted (Figure S7A-C). Only proteins with normalized H/L ratio > 1.0 in both replicates were defined as enriched, yielding a list of 268 proteins in the RinID nuclear proteome (Figure S7D, Table S7). Gene Ontology analysis reveals that 82% (219 out of 268) of proteins in the list have prior nuclear annotations, which is comparable to the specificity of TurboID nuclear proteome (79%)^8^ and substantially higher than the percentage in the human proteome (36%) (Figure 4A, 4B). The lower coverage of RinID may result from the use of histone protein H2B as the bait for proximity labeling. In contrast, TurboID was targeted to the nucleoplasm via fusion with a nuclear localization sequence. Thus, the proteome identified by RinID is more focused on histone and chromtain-associated proteins, but contains less other nuclear proteins. For example, among the top 20 nuclear proteins enriched by H2B RinID, 12 (60%) have chromosome/chromatin annotations in Gene Ontology. The coverage of RinID nuclear proteome is much higher than a recently reported proximity labeling method using chromatin-targeted small molecule photocatalyst dibromofluorophore-Hoechst, in which only 10 nuclear proteins were identified^23^.

**Figure 4.**
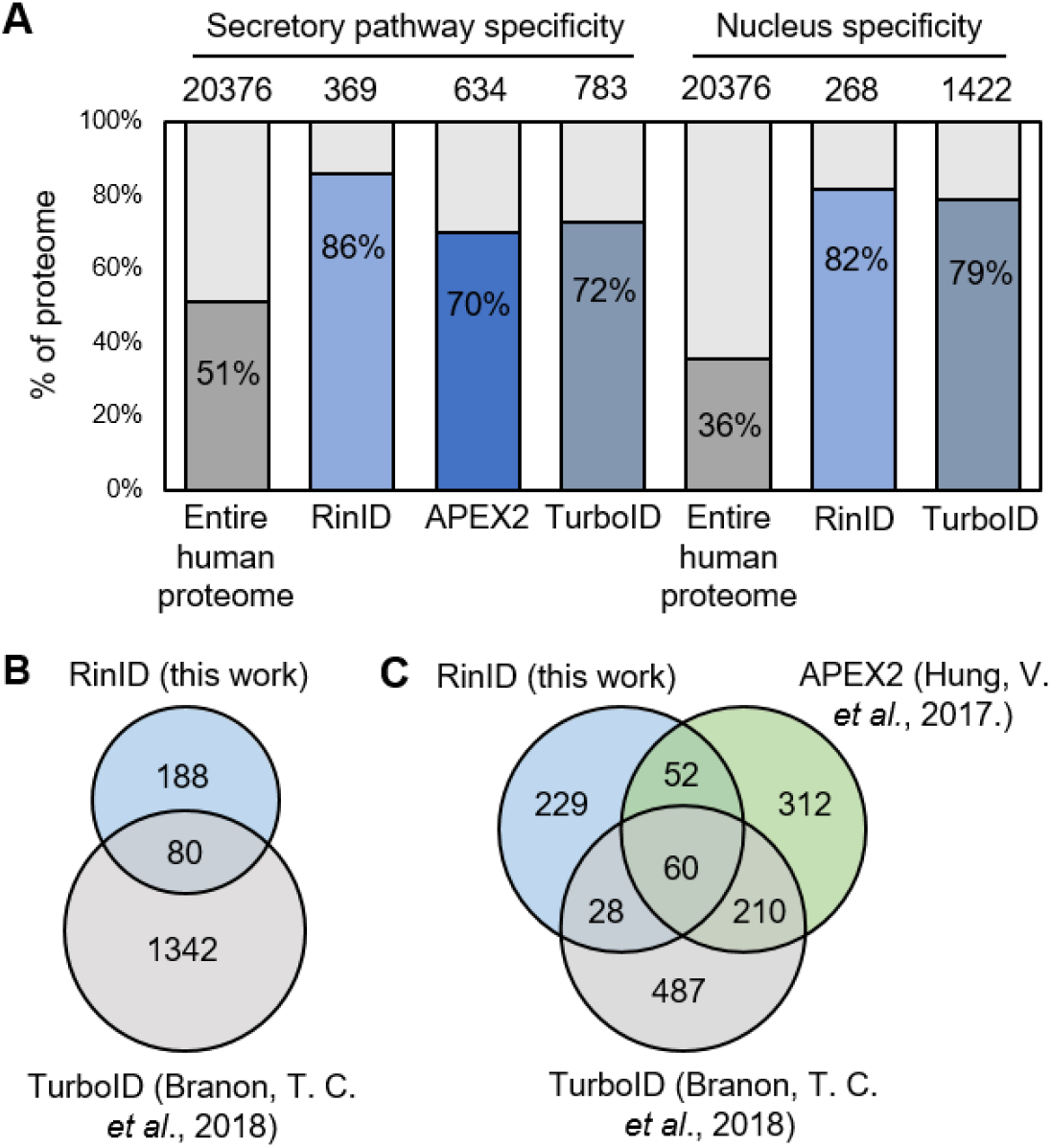
Subcellular proteomic profiling with RinID in the nucleus and ER membrane (ERM). **A)** Comparison of spatial specificity of proteomic data derived from RinID and other proximity labeling methods at ERM and nucleus. **B-C)** Comparisons of proteomic coverage by RinID and other proximity labeling methods at the nucleus (B) and ERM (C).

For the profiling of ERM proteome, HEK293T cells expressing ERM-targeted SEC61B-miniSOG fusion protein were labeled with 20 mM PA for 15 min in two biological replicates, with blue light omission as the negative control (Figure S8A-B). Proteins with normalized H/L ratio over 1.0 in both replicates were defined as enriched proteins (Figure S8C, Table S8). Among the list of 369 enriched proteins in the ERM RinID proteome, 86% (318 out of 369) are annotated as secretory pathway proteins, which is higher in specificity when compared to TurboID (72%)^8^ and APEX2 (70%)^10^ (Figure 4A, 4C). In comparison, only 51% of the human proteome are secretory pathway proteins (Figure 4A). Admittedly, the coverage of RinID datasets in both compartments are lower that previously reported TurboID and APEX2 datasets, suggesting that the labeling sensitivity needs to be further improved. Nevertheless, that RinID is capable of identifying proteins not previously covered by TurboID and APEX2 indicates that this new method could complement existing techniques by expanding the overall coverage.

The strong dependence of RinID on light illumination could be leveraged to achieve pulse-chase protein labeling (Figure 5A). While long-term and intense irradiation of miniSOG could lead to excessive protein oxidation and even cell death^24^, it might be possible to balance between efficient protein labeling and low cytotoxicity by carefully tuning the dosage of blue light. To reduce illumination time, we chose an engineered miniSOG variant, SOPP3, with improved singlet oxygen quantum yield^25^.

**Figure 5.**
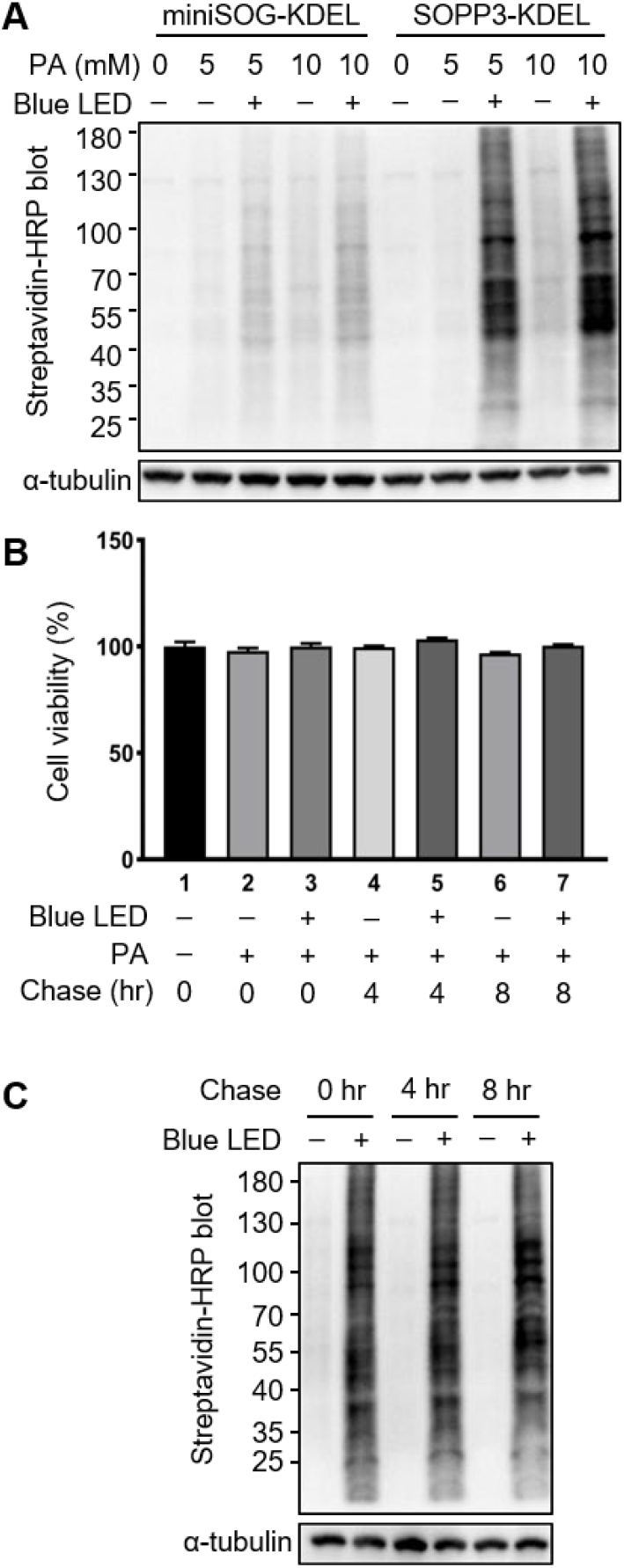
Pulse-chase labeling of secretory pathway proteome with RinID. **A)** Western blot of ER lumen proteome labeled with miniSOG or SOPP3 at different PA concentrations, with blue light illumination at 30 mW·cm^-2^ for 5 min. **B-C)** Cell viability assay (B) and Western blot analysis (C) of Hela cells at 0, 4, and 8 hours after SOPP3 labeling in the ER lumen with 5 min blue light irradiation and 5 mM PA.

We constructed HEK293T and Hela cell lines stably expressing SOPP3 targeted to the mitochondrial matrix and ER lumen. Western blot and immunofluorescence imaging analysis revealed stronger labeling intensity by SOPP3 over miniSOG with 5 min blue light illumination and 5 mM PA (Figure 5A, S9). Thereafter, labeled Hela cells were “chased” by culturing in the absence of PA probe and blue light for another 4 – 8 hours. Mitochondrial activity assay showed no changes in cell viability during the 8 hours chase period (Figure 5B). Western blot analysis also showed similar pattern of protein biotinylation over the course of chase (Figure 5C). Collectively, these results indicate that pulse labeling with SOPP3 causes minimal cytotoxicity and is thus applicable for pulse-chase experimental schemes, which is useful for future investigations of subcellular protein dynamics (e.g. intracellular trafficking, metabolic turnover, post translation modification) upon external stimulus.

## DISCUSSION

In summary, we have developed RinID, a light-activated proximity-dependent protein labeling method for profiling subcellular proteome. We characterized its labeling mechanism and identified amino acid residue of photo-oxidation. Using two cell lines and multiple subcellular organelles (i.e. mitochondria, nucleus and ER), we demonstrated the exceptional spatial specificity of RinID by both fluorescence imaging and quantitative proteomics.

Over the past decade, engineered peroxidases (e.g. APEX^5^, APEX2^6^, HRP^26^) and biotin ligases (e.g. BioID^7^, TurboID^8^) have been employed as powerful tools for proximity labeling. The spatial specificity and coverage of RinID is comparable to these methods. Meanwhile, the difference in amino acid preference suggests that RinID may complement existing techniques to improve the overall coverage. Compared to APEX, RinID is less toxic by avoiding the use of H2O2. Compared to TurboID, the light-triggered RinID labeling offers better temporal control of the reaction, which is more suitable for studying dynamic changes in the subcellular proteome. It should be noted that miniSOG has been previously used for probing protein-protein interactions by photo-oxidizing the cysteine residues of nearby proteins, which form disulfide bonds with a biotin-conjuated thiol probe^27^. However, the low efficiency of forming disulfie bond in the reducing cytosolic environment has hampered its applications for proteome-wide identification.

While RinID has proven efficient in multiple subcellular organelles, a few problems still need to be solved in the future. First, the background generated by PA probe needs to be deducted by quantitative MS experimental design, which may complicate data analysis. Second, the 15 min labeling time is still longer compared to APEX2. Third, the tissue penetration of blue light is typically restricted to <1 mm, thus limiting applications to tissue samples such as brain slice. Similar to other proximity labeling methods, these problems could be solved by the development of better probes with higher specificity and labeling efficiency, and by the design of photocatalysts with higher quantum yield of singlet oxygen and red-shifted excitation spectrum. With further development of such probes and photocatalysts, RinID would become a powerful tool for high spatiotemporal resolution identification of subcellular proteomes.

## AUTHOR CONTRIBUTIONS

P.Z. conceived the project. F.Z. and P.Z. designed experiments. F.Z. and C.Y., and X.Z. performed all experiments. F.Z. and P.Z. analyzed data and wrote the paper with inputs from other authors.

## ACKNOWLEDGMENT

We thank Y. Li for advice on MS sample preparation, L. Peng for assistance with image analysis, Y. Fu, Y. Zhou, and G. Wang for assistance with probe synthesis, all lab members for helpful discussions. This work was supported by the Ministry of Science and Technology (2018YFA0507600, 2017YFA0503600), the National Natural Science Foundation of China (32088101, 21727806). PZ is sponsored by Bayer Investigator Award. The measurement of NMR was performed at the Analytical Instrumentation Center of Peking University. We thank the Analytical Instrumentation Center in Peking University for assistance with MS sample identification and Ms. W. Zhou for help with MS results analysis.

